# Empowering conservation practice with efficient and economical genotyping from poor quality samples

**DOI:** 10.1101/349472

**Authors:** Meghana Natesh, Ryan W. Taylor, Nathan Truelove, Elizabeth A. Hadly, Stephen Palumbi, Dmitri Petrov, Uma Ramakrishnan

**Affiliations:** National Centre for Biological Sciences, TIFR, Bellary Road, Bangalore - 560065, India; Sastra University, Tirumalaisamudram, Thanjavur – 613401, India; Department of Biology, Stanford University, Stanford, CA - 94305, USA; End2End Genomics LLC, Davis, CA; Hopkins Marine Station, Stanford University, Pacific Grove, CA – 93950, USA

**Author notes:** Equal contribution (Meghana Natesh, Ryan W. Taylor, Dmitri Petrov, Uma Ramakrishnan). Correspondence: Dmitri Petrov, Department of Biology, 371 Serra St., Stanford University Stanford, CA 94305-5020, USA, and Uma Ramakrishnan, Senior Fellow, Wellcome Trust/DBT India Alliance, NCBS TIFR, Bangalore, 560065, India.

**Keywords:** conch, conservation genetics, endangered species monitoring, genotyping, multiplex PCR, non-invasive samples, SNPs, tigers

## Abstract

1. Moderate to high density genotyping (100+ SNPs) is widely used to determine and measure individual identity, relatedness, fitness, population structure and migration in wild populations.
2. However, these important tools are difficult to apply when high-quality genetic material is unavailable. Most genomic tools are developed for high quality DNA sources from labor medical settings. As a result, most genetic data from market or field settings is limited to easily amplified mitochondrial DNA or a few microsatellites.
3. To enable genotyping in conservation contexts, we used next-generation sequencing of multiplex PCR products from very low-quality DNA extracted from feces, hair, and cooked samples. We demonstrated utility and wide-ranging potential application in endangered wild tigers and tracking commercial trade in Caribbean queen conch.
4. We genotyped 100 SNPs from degraded tiger samples to identify individuals, discern close relatives, and detect population differentiation. Co-occurring carnivores do not amplify (e.g. Indian wild dog/Dhole) or are monomorphic (e.g. leopard). 62 SNPs from conch fritters and field-collected samples were used to test relatedness and detect population structure.
5. We provide proof-of-concept for a rapid, simple, cost-effective, and scalable method (for both samples and number of loci), a framework that can be applied to other conservation scenarios previously limited by low quality DNA samples. These approaches provide a critical advance for wildlife monitoring and forensics, open the door to field-ready testing, and will strengthen the use of science in policy decisions and wildlife trade.

## Introduction

Stemming the tide of global species decline requires continuous monitoring and nimble, adaptive management to promote species recovery. Effective monitoring relies on identifying species presence and the ability to track specific individuals and their familial relationships. While species recovery is critically dependent on tracking individuals and their dynamics locally, integrating data across the species range allows monitoring of global large-scale threats including population range reduction and illegal wildlife trade.

In principle, all of these goals can be achieved via genotyping a modest number of loci such as microsatellites or single nucleotide polymorphisms (SNPs). To study endangered species that are rare and elusive, approaches must be able to accommodate non-invasive sources of DNA such as feces, shell, feathers, hair, and saliva, which yield impure, mixed, and/or extremely small amounts of degraded DNA. Moreover, market samples generated by wildlife trade may be processed, cooked, dried, or mixed with other species, again providing low quality and often mixed DNA. Current approaches tend to require relatively large amounts of DNA (Carroll et al. 2018, nanograms of DNA) from the target species, or demand expensive and generally inefficient enrichment strategies (Chiou & Bergey, 2018). Approaches designed for lower concentration DNA samples (Kraus et al. 2015) require expensive and specialized equipment.

Here we demonstrate that a multiplex PCR approach followed by next-generation sequencing satisfies all the requirements necessary for inexpensive, fast, and easy genotyping of low-quality samples. Our approach is similar to GT-seq (Campbell, Harmon, & Narum, 2014) but can use publicly available software for designing primers and calling SNPs, and targets only very short fragments in order to succeed with degraded DNA. We illustrate the power of this method for two endangered species in very divergent conservation contexts and real-life settings: genotypes from feces, shed hair, and saliva found on killed prey from wild Indian tigers and from CITES-regulated Caribbean queen conch imported to the US and sold in fried fritters (method schematic Fig. 1a). Methods include DNA extraction, a multiplex PCR, a second barcoding PCR, Illumina miseq sequencing and bioinformatics for SNP genotyping.

**Figure 1a.**
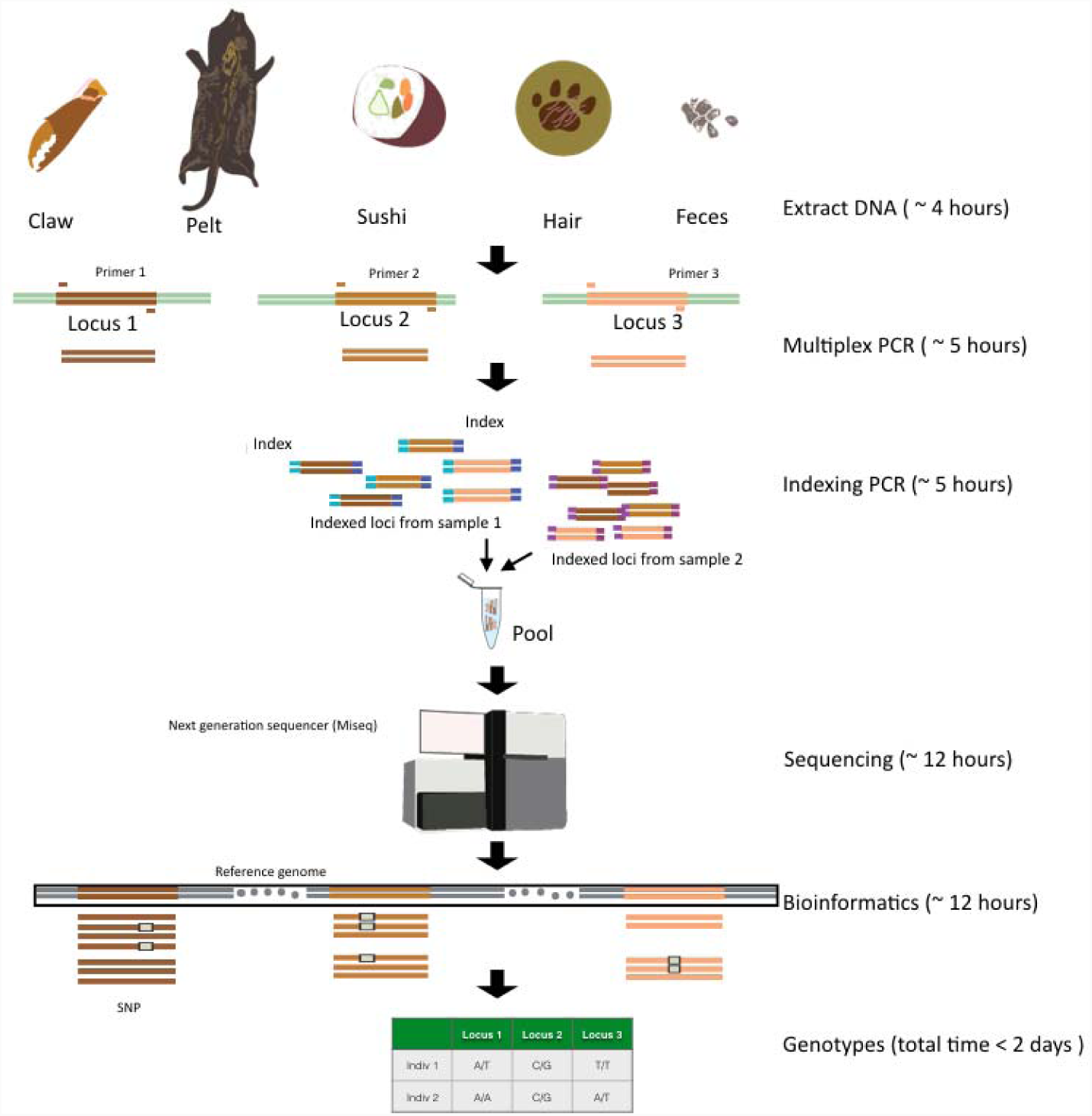
A schematic for protocol and approximate time taken. Available samples include a variety of sources. Details in the main and supplementary text.

**Figure 1b.**
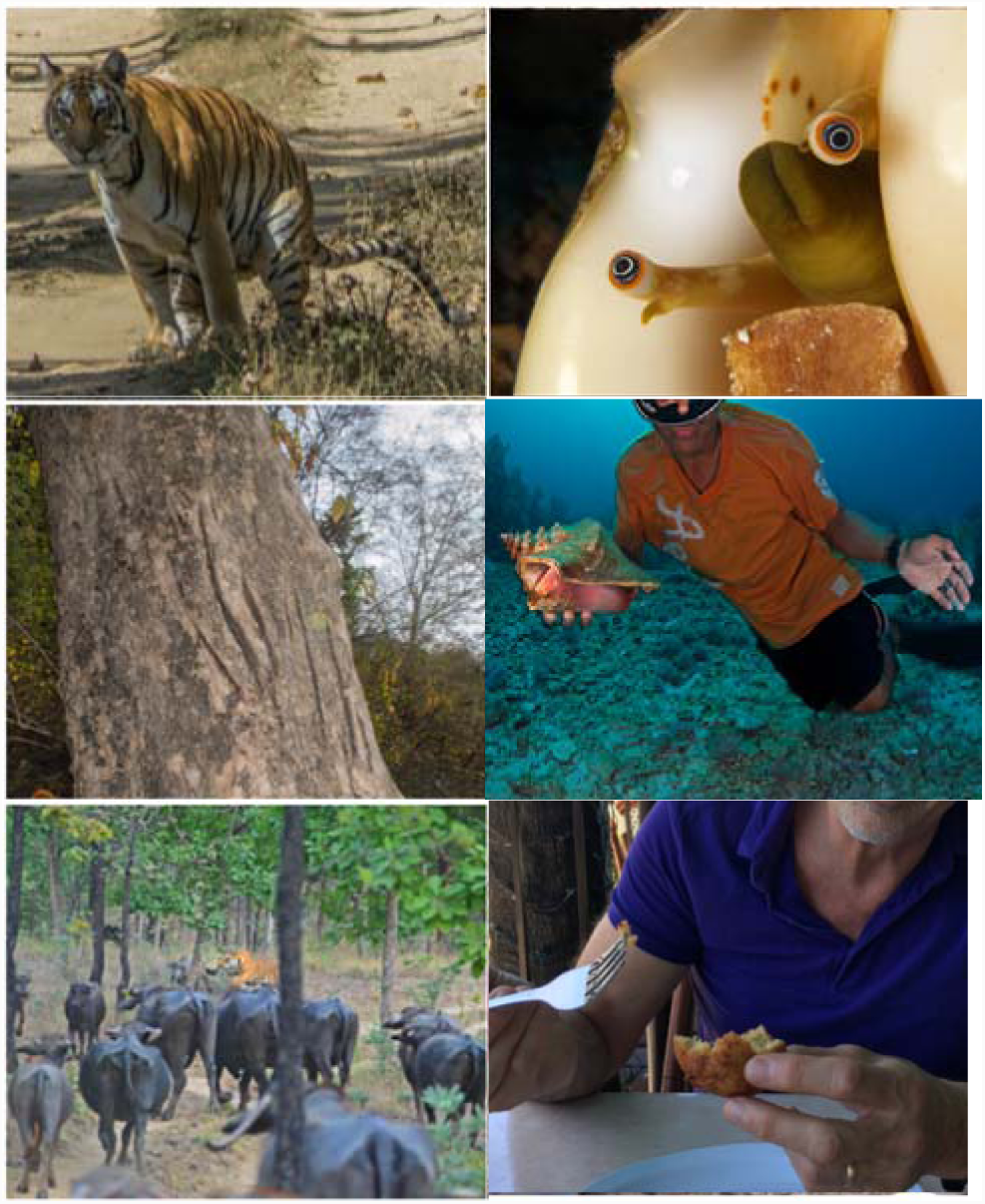
On the left: a tiger defecating (photo: Himanushu Chhattani), a scrape mark (and associated hair samples, photo: Kaushal Patel), and a tiger hunting (photo: Shantanu Prasad, associated lick marks). On the right, a conch emerging from its shell, a fisherman holding his catch and a fritter.

The tiger (*Panthera tigris*), a charismatic carnivore classified as endangered by the IUCN red list. Their distribution across 14 countries (Goodrich et al. 2015) makes it critical that locally collected data be comparable across their range. Genotyping of scat or hair, along with rapid forensic testing of confiscated skins or other traded parts can verify species, individual identity, and source populations. We developed a multiplex primer set for 192 SNP loci and tested them on fecal, tissue, saliva, and hair samples from captive and wild tigers. We also genotyped two sympatric carnivores, the Dhole (*Cuon alpinus*), and the leopard (*Panthera pardus*), that may be confused with tigers when targeting noninvasive samples.

Our second example illustrates the use of this approach even when reference genomes are unavailable, and again highlights use in difficult samples: in this case fried conch fritters. Although formerly abundant, the queen conch (*Strombus gigas*) was listed in CITES Appendix II in 1990, which allows for control of trade to reduce over-exploitation. Identifying geographic ancestry of illegally traded queen conch products in Florida markets will aid conservation action that will allow recovery of this formerly lucrative fishery. Because the most direct access to imported conch is from the hundreds of restaurants in Florida, we sought techniques that would allow genotyping from the most abundant menu item, fried conch fritters.

## Methods

### Sample Collection

For tigers, multiple samples from 13 captive tigers (USA zoos) representing different scenarios were used for standardization (details in Table S1). Blood and corresponding fecal swabs from captive tigers, (including one parent-offspring and one sibling pair) were collected in India. Wild tigers were sampled noninvasively from multiple protected areas across India (Table S2).

Scat and saliva (from predator bites on the prey) were sampled by swabbing the surface of the sample with moistened synthetic swabs (Ramón-Laca et al. 2015); tips were stored in lysis buffer in 2ml microcentrifuge. Hair was sampled using forceps and stored in ziplock bags. DNA from legacy fecal samples (collected in alcohol in 2014) were also tested. DNA extraction (first step in Figure 1a) and quantification are described in SM1. Two leopards and one Dhole were included. Most samples were genotyped in triplicate.

Tissue samples from live caught Queen Conch mantle were collected using a sterilized biopsy forceps were preserved in RNALater. Samples from conch fritters purchased from Miami restaurants were frozen until dissected to isolate animal tissue fragments (SM 3a).

### SNP Identification and filtering

#### Tigers

We identified SNPs from whole genome sequencing of 75 tigers of wild and captive origin from *P. t. tigris, P. t. jacksoni, P. t. altaica, and P. t. sumatrae* subspecies (SM 1a). Alignment, filtering and pruning were used to identify SNPs (SM 1a). We calculated minor allele frequencies (MAF) for each SNP within each subspecies, and retained SNPs with MAF 10, 15, 20, 25, and 30 percent. We identified fixed SNPs that differentiated populations. We prioritized SNPs within contigs greater than 10, 5, and 1MB respectively. We selected 10 differentiating SNPs from each subpopulation, and 9992 polymorphic SNPs from each of the aforementioned MAF cutoffs, for a total of 50,000 SNPs.

#### Conch

We extracted RNA from 96 *L. gigas* individuals (SM 3a, table S3). Four queen conch individuals (one each from Aruba, Belize, Florida, and St. Eustatius) were imported into TRINITY v2.2.0 (Grabherr et al. 2013) to assemble a *de novo* transcriptome (SM 3a).480,962 SNPs were discovered by aligning 96 conch sequences from 6 populations in the Caribbean to the assembled transcriptome(SFG pipeline -https://github.com/bethsheets/Population-Genomics-via-RNAseq, SM 3a).

### Primer Design

Publically available Primer3(amplicon size 50-90 bp, primer size 17-25 bp, and T^m^ of 60-61ºC) was used to design primers for tiger SNPs. Conch primers were designed using the G4C (Genotyping for Conservation) script library and the same criteria. No attempt was made to identify incompatibilities between primer-pairs and 192 primer pairs were shortlisted for both species (SM 1b, SM 3b).GT-seq indexes and adapters were used (Campbell, Harmon, & Narum, 2014).

### PCR amplification and sequencing

Library preparation consisted of an initial multiplex PCR reaction, a second PCR reaction to add sequencing adapters and indexes, and sample pooling. Sample input DNA volume was adjusted to a maximum of 1ng per reaction. The multiplex PCR simultaneously amplified all target regions for each sample separately (in a 96 well plate). The second PCR reaction added a combination of forward (i5) and reverse (i7) Illumina indexes to uniquely identify each sample. The sequencing library contained equal volumes of each sample’s barcoded product and was cleaned with Ampure beads. Sequencing of single 50bp reads was performed on Illumina MiSeq. A detailed protocol is in Supplement 1f. Alignment and genotype calling followed GT-Seq for the conch study (SM3b; Campbell, Harmon, & Narum, 2014) and used standard open source tools for the tiger study (SM1h). Figure 1a illustrates the steps described above.

For tigers, genotyping success, genotype concordance across replicates, relatedness between pairs of individuals of known and unknown relationship, probability of identity of the SNP panel and population structure were assessed (details in SM1h). For conch, genotyping success and genotype concordance were tested by comparing SNPs across replicate conches. Genetic distance and ability to assign the conch samples to the correct population was estimated by comparison of the 96 transcriptome samples (details in SM 4).

## Results

### Tiger

126 targets (66% of 192 attempted) produced the most consistent results (SM2b, Figure S1, Table S4), and were then tested on non-invasive samples from wild tigers across India and zoo individuals (Fig. 1b, Table S2).

The 126 SNP panel for wild tigers had a high overall genotyping success rate (Fig. 2, SM 2c). An average of 95 SNPs (75%, range: 4 - 114) were successfully typed across all samples. When tiger DNA was > 0.01ng/ul (all sample types) an average of 105 SNPs (83%, range: 48 - 114) were typed.

**Figure 2.**
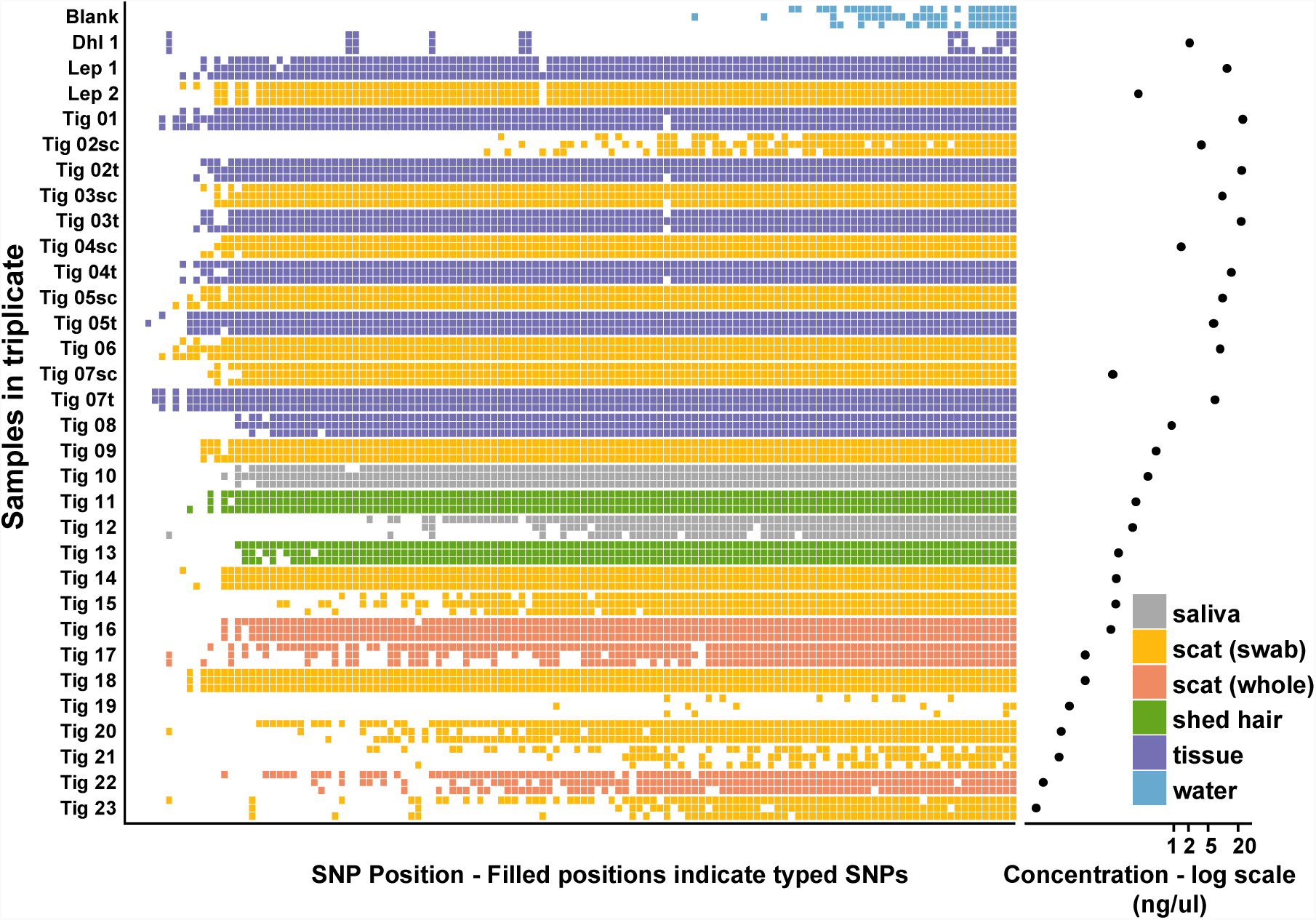
SNP typing success (in triplicate) for various tiger samples and controls. Filled cells indicate successfully typed SNPs; color indicates sample type. DNA concentrations are on the right.

Sample replicates were highly concordant with a high proportion of genotype matches (n=28 triplicates, genotype concordance, mean: 0.957, range: 0.757 - 1.0). Different sample types from the same individual also had highly concordant genotypes (n=5, concordance, mean 0.97; range 0.91 - 0.99). Our error rates were comparable to low microsatellite genotyping error rates in some studies (Thaden et al. 2017) or lower other non-invasive studies (Mondol et al. 2014). Our probability of misidentification was vanishingly low (pID_sibs_ = 1.6E-22).

The co-occurring carnivores, leopard and dhole, could be distinguished from tiger genotypes. The dhole tissue sample had very poor amplification success (mean across replicates <10 SNPs) as expected. Leopards had high amplification success (mean: 110 SNPs), but SNPs were monomorphic and nearly identical across two individuals (mean: 0.03% mismatches).

The known parent-offspring and sibling pairs (captive individuals, India) had relatedness values close to expected values (0.5, Figure 3). Observed pair-wise relatedness was higher within than between known genetic clusters or populations (see Natesh et al. 2017). As expected, relatedness among individuals from a small, isolated population (NW, Figure 3) was high. Wild individuals fell into three genetic clusters/populations as expected (Figure S2).

**Figure 3:**
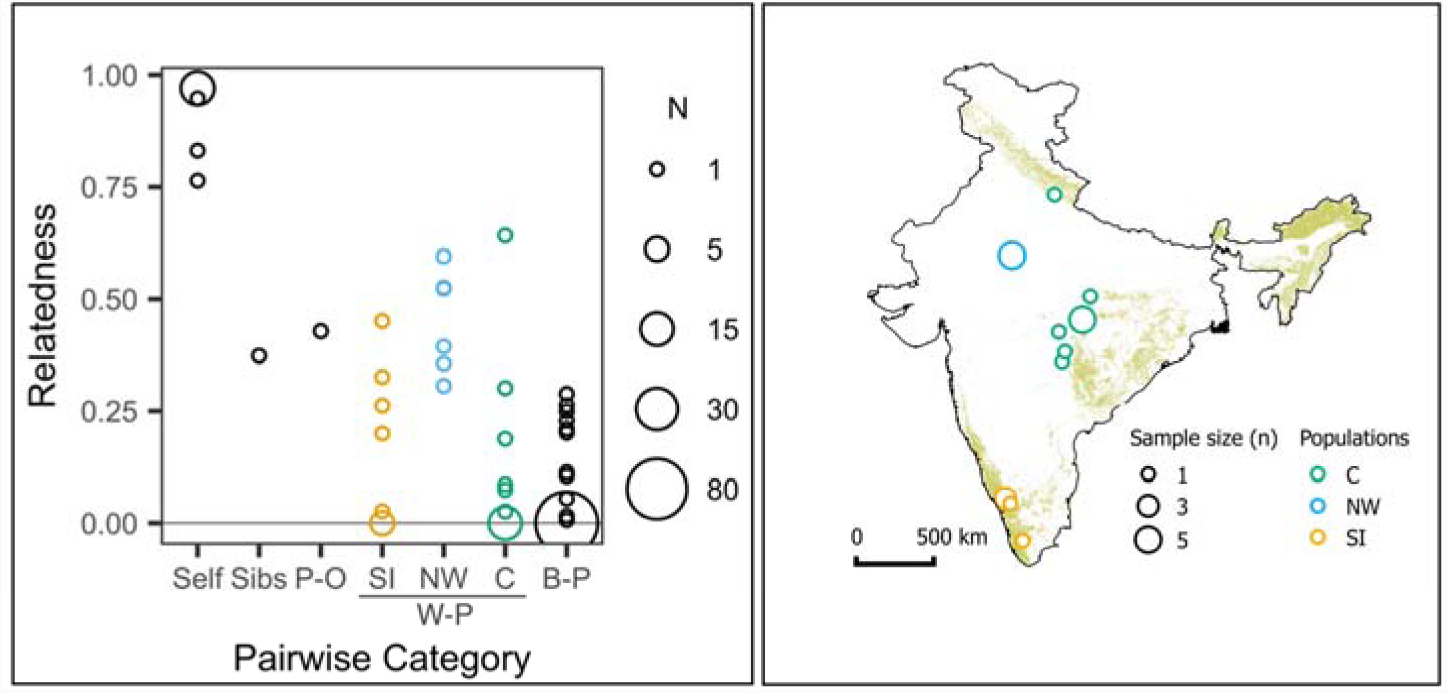
Pairwise relatedness (PI_HAT): Self: three replicates of each sample; same individual but different sample, Sibs: siblings, P-O: parent-offspring pair, W-P: within population, B-P: between populations. True relatedness unknown for W-P and B-P. Sampling locations and corresponding colours represented in the India map on right.

### Queen Conch

We tested a 192 primer-pair panel on 279 conch samples from 14 populations, including 48 fried conch fritters from Miami, Florida restaurants. Our SNP success was lower for conch than tigers (SM 4). Approximately half the 192 conch primer-pairs failed to provide data, but 62 targets reached an 86% success rate similar to the 126 good tiger targets (Fig. S3). Replicate samples shared 99%-100% of their alleles, and different processed conch samples were genetically identical, suggesting recapture. There were no obvious close relatives among the samples (Fig. S4 and S5). Outlying islands (Aruba, St. Eustatius) were genetically differentiated from the central Caribbean and Florida (F_ST_ = 0.037, 0.048 respectively, Table S5). Samples from the same island group (e.g. Florida, Bahamas) or the same coast were not differentiated. These patterns parallel a recent survey of queen conch with microsatellites (Truelove et al. 2017).

Conch DNA from deep-fried fritters had lower success rates than from biopsies. However, success was high enough for individual identification and initial population comparison. Twenty-three of the fritters were probably not Queen conch (fewer than 18 SNPs amplified), 8 fritters revealed poor SNP amplification (average 41% success), and 17 fritters were comparable to fresh samples (77% vs 86% success, Fig. S3). These samples revealed lower (average) genetic identity to Florida populations (average 78.6 – 80.5 alleles shared, Fig. S6) compared to Puerto Rico, or Andros Island (81.2-82.3, Fig. S6). While positive population identification may require greater geographic sampling and more SNPs (from several whole genomes), our pilot data suggest that the fritters we genotyped are less likely to be from the Florida Keys and Nassau. Importantly, high resolution nuclear data was readily obtained from processed commercial samples to address key conservation challenges.

## Discussion

Our pilot datasets provide proof of concept for multiplex SNP genotyping of non-invasive and processed market samples from two species with vastly different physiologies, ecologies, and conservation challenges. The approach is successful for degraded, cooked, mixed, or small and low-concentration samples (down to 10^-3^ ng DNA/ul), making it an ideal tool for monitoring individuals under field conditions or from commercial markets.

Conservation practitioners assume that genetics is expensive. However, our method is cheap, while providing rich information important for conservation. Designing a similar protocol for a new species of interest would include costs for method development and implementation. Development costs include polymorphic SNP ascertainment, primer design and synthesis. However, note that SNPs have already been identified for many endangered and fisheries species (e.g. Steiner et al. 2013). If no SNPs have been identified, practitioners could ascertain SNPs using whole genome sequencing (e.g. genome assembly of reasonable quality ∼$2,000, Armstrong et al., 2017) and pooled sequencing of 10 individuals at approximately 50X total coverage, ∼ $1000). The upfront cost of primer design and synthesis is between $1,000 to $10,000 (for 100 to 1,000 primer pairs, $10 per primer pair). Once synthesized, primers can be used for 38,000 reactions (∼ 400 plates). The continual advance in sequencing and oligo synthesis will drive down these initial development costs. Most important and attractive to conservationists, we estimate implementation costs (for 1000 SNPs) can be as low as $5 per sample (when processing several hundred samples).

Increasing the number of SNPs beyond a few hundred can provide additional information. For this pilot, we constrained the number of targeted SNPs, but it should be possible to target many more. Primers chosen to amplify clusters of closely located SNPs should allow detection of very recent inbreeding using long contiguous runs of homozygosity (Kirin et al. 2010). Linked SNPs could generate microhaplotypes, particularly useful in pedigree reconstruction (Baetscher et al. 2018). SNP panels could allow simultaneous species, individual, and diet identification for sets of species, e.g. large carnivores and common prey species in India or sub-Saharan Africa.

Effective monitoring of individuals, populations and species is critical to designing rapid conservation action and management of endangered species like tigers. Small, isolated populations will require inbreeding management, genetics-based population assessment, and genetically-informed introduction strategies. Likewise, identification of commercial products from illegal fishing, bush meat hunting or highly processed market samples provides important management information. Ability to assay such samples could provide a powerful incentive to enforce local conservation laws. Rapid genetic monitoring of endangered species from commonly occurring non-invasive samples will provide a data-pathway to species recovery. We believe that multiplex PCR presents an example of such rapid, accessible, cheap and efficient technology that will make this possible.

## Acknowledgements

San Diego Zoo Global, CSC at Oakland Zoo, SF Zoo, Bronx Zoo, WCS Russia, El Paso Zoo, Performing Animal Welfare Society, WCS India, CWS India, WCT, Bannerghatta Zoo, Himanshu, Anubhab, Prachi, Aditya, Arun, Erica Calcagno, Jackie Gai for tiger samples/data, and forest departments of Rajasthan, Karnataka, Madhya Pradesh and American Zoo Association, Tiger SSP for permissions. Awadhesh Pandit and NCBS NGS facility. Steve Box, and Steve Canty.

## Data Availability

Raw sequences are uploaded to NCBI (PRJNA516037) and primer sequences for both tigers and conch are in Supplementary File (Primer_Sequences).

## References

Armstrong, E. E., Taylor, R. W., Prost, S., Blinston, P., Der, E. Van, Madzikanda, H., …Petrov, D. (2017). Entering the era of conservation genomics: Cost-effective assembly of the African wild dog genome using linked long reads. BioRxiv. doi:http://dx.doi.org/10.1101/195180.

Baetscher, D. S., Clemento, A. J., Ng, T. C., Anderson, C., & Garza, J. C. (2018). Microhaplotypes provide increased power from short-read DNA sequences for relationship inference. Molecular Ecology Resources, 18(November 2017), 296–305. doi:10.1111/1755-0998.12737

Campbell, N. R., Harmon, S., & Narum, S. R. (2014). Genotyping-in-Thousands by sequencing (GT-seq): A cost effective SNP genotyping method based on custom amplicon sequencing. Molecular Ecology Resources, n/a-n/a. doi:10.1111/1755-0998.123571

Carroll, E. L., Bruford, M. W., Dewoody, J. A., Leroy, G., Strand, A., Waits, L., & Wang, J. (2018). Genetic and genomic monitoring with minimally invasive sampling methods. Evolutionary Applications, 1–26. doi:10.1111/eva.12600

Chiou, K. L., & Bergey, C. M. (2018). Methylation-based enrichment facilitates low-cost, noninvasive genomic scale sequencing of populations from feces. Scientific Reports (January), 1–10. doi:10.1038/s41598-018-20427-9

Fitak, R. R., Naidu, A., Thompson, R. W., & Culver, M. (2016). A New Panel of SNP Markers for the Individual Identification of North American Pumas. Journal of Fish and Wildlife Management, 7(1), 13–27. doi:10.3996/112014-JFWM-080

Goodrich, J. M., Lyam, A., Miquelle, D. G., Wibisono, H. T., Kawanishi, K., Pattanavibool, A., … Karanth, U. K. (2015). Panthera tigris. The IUCN Red List of Threatened Species 2015. Iucn, 8235. doi:e.T15955A50659951. http://dx.doi.org/10.2305/IUCN.UK.2015-2.RLTS.T15955A50659951.en.

Grabherr, M. G., Haas, B. J., Yassour, M., Levin, J. Z., Thompson, D. A., Amit, I., … Regev, A. (2013). Trinity: reconstructing a full-length transcriptome without a genome from RNA-Seq data. Nature Biotechnology, 29(7), 644–652. doi:10.1038/nbt.1883.Trinity

Kirin, M., Mcquillan, R., Franklin, C. S., Campbell, H., Mckeigue, P. M., & James, F. (2010). Genomic Runs of Homozygosity Record Population History and Consanguinity. PLoS ONE, 5(11), 1–7. doi:10.1371/journal.pone.0013996

Kraus, R. H. S., vonHoldt, B., Cocchiararo, B., Harms, V., Bayerl, H., Kühn, R., … Nowak, C. (2015). A single-nucleotide polymorphism-based approach for rapid and cost-effective genetic wolf monitoring in Europe based on noninvasively collected samples. Molecular Ecology Resources, 15(2), 295–305. doi:10.1111/1755-0998.12307

Mondol, S., Kumar, N. S., Gopalaswamy, A., Sunagar, K., Karanth, K. U., & Ramakrishnan, U. (2014). Identifying species, sex and individual tigers and leopards in the Malenad-Mysore Tiger Landscape, Western Ghats, India. Conservation Genetics Resources, 353–361. doi:10.1007/s12686-014-0371-9

Natesh, M., Alta, G., Nigam, P., Jhala, Y. V, Zachariah, A., Borthakur, U., … Ramakrishnan, U. (2017). Conservation priorities for endangered Indian tigers through a genomic lens. Scientific Reports, 7(1), 1–11. doi:10.1038/s41598-017-09748-3

Ramón-Laca, A., Soriano, L., Gleeson, D., & Godoy, J. A. (2015). A simple and effective method for obtaining mammal DNA from faeces. Wildlife Biology, 21(4), 195–203. doi:10.2981/wlb.00096

Snyder-Mackler, N., Majoros, W. H., Yuan, M. L., Shaver, A. O., Gordon, J. B., Kopp, G. H., … Tung, J. (2016). Efficient Genome-Wide Sequencing and Low Coverage Pedigree Analysis from Non-invasively Collected Samples. Genetics, 203(June), 699–714. doi:10.1534/genetics.116.187492

Steiner, C. C., Putnam, A. S., Hoeck, P. E. A., & Ryder, O. A. (2013). Conservation genomics of threatened animal species. Annual Review of Animal Biosciences, 1, 261–281. doi:10.1146/annurev-animal-031412-103636

Thaden, A. Von, Cocchiararo, B., Jarausch, A., & Jüngling, H. (2017). Assessing SNP genotyping of noninvasively collected wildlife samples using microfluidic arrays, (May), 1–13. doi:10.1038/s41598-017-10647-w

Truelove, N. K., Box, S. J., Aiken, K. A., Boman, E. M., Booker, C. J., Byfield, T. T., … Stoner, A. W. (2017). Isolation by oceanic distance and spatial genetic structure in an overharvested international fishery, 1292–1300. doi:10.1111/ddi.12626

