## Supplementary information for "Empowering conservation practice with efficient and economical genotyping from poor quality samples"

**This PDF file includes:**

Supplementary Materials, Methods and Results

1. Tiger genotyping: Materials and Methods
   a. Primer design
   b. Lab protocol
   c. Samples chosen for testing
   d. Bioinformatics and analysis
2. Tiger genotyping: Supplemental Results

a. The effect of sample DNA concentration on genotype success

b. 192 SNP panel

c. 126 SNP panel

3. Conch Genotyping: Materials and Methods
   a. Conch transcriptome
   b. SNP primer design and testing
4. Conch Genotyping: Supplemental Results

Supplementary Figures S1: S6

Supplementary Tables S1: S5

1. **Tiger genotyping: Materials and Methods**

1a. Primer design

*SNP Identification and filtering*

We identified single nucleotide polymorphisms from whole genome sequencing of 75tigers of wild and captive origin from *P. t. tigris, P. t. jacksoni, P. t. altaica, and P. t. sumatrae* subspecies. Full details about the samples and whole genome sequencing are in preparation (Armstrong et al. Unpublished Manuscript). To identify polymorphic sites for this genotyping experiment we aligned the WGS reads with bwa mem (v0.7.15-r1140; Li & Durbin 2009) to a 10x Genomics based reference genome (Armstrong et al. Unpublished Manuscript), and marked duplicates with the Picard Tools MarkDuplicates (v2.9.0-1-gf5b9f50-SNAPSHOT; Broad Institute) command. We called SNPs with FreeBayes (v1.1.0-3-g961e5f3-dirty;Garrison & Marth, 2012) with the '--use-best-n-alleles 4' option specified. Finally we filtered the variants for biallelic SNPs with a site wide quality score of 30, and individual genotype quality scores of 30.We then pruned the filtered SNPs for linkage disequilibrium using PLINK (v1.90b4.6indep; Purcell et al., 2007) with a window size of 50 variants, a shift of 5 variants, and a variance inflation factor of 2. To identify a pool of potential target SNPs we calculated minor allele frequencies (MAF) for each SNP for each subspecies. We then identified SNPs with minor alleles present at a minimum frequency of 10, 15, 20, 25, and 30 percent. We also identified differentiating SNPs that were fixed in all populations and the minor allele only existed in one population. We selected 50,000 SNPs to design primers for. We prioritized SNPs with contigsize greater than 10, 5, and 1MB respectively. We selected 10 differentiating SNPs from each subpopulation, and 9992 polymorphic SNPs from each of the aforementioned MAF cutoffs, for a total of 50,000 SNPs.

*Primer Design*

We used Primer3 to design primers for these 50,000 SNPs using the following parameters: PRIMER_OPT_SIZE=20, PRIMER_MIN_TM=60, PRIMER_MAX_TM=61, PRIMER_MIN_SIZE=17, PRIMER_MAX_SIZE=25, PRIMER_MAX_NS_ACCEPTED=0, PRIMER_PRODUCT_SIZE_RANGE=50-90. From the resulting SNPs for which Primer3 was able to design primer pairs, we selected the 17 differentiating SNPs for which primers were found, and randomly selected 35 SNPs with MAF of 10, 15, 20, 25, and 30 percent, for a total of 192 SNPs.We did not perform any compatibility checks between the primer pairs, or any mispriming or off-target amplification checks. A more thorough interrogation of likely primer success would probably reduce the number of failed primers, but as we show here it is not necessary.

We used GT-seq(Campbell, Harmon, & Narum, 2014) adapters and indexes.

1b. Lab protocol

*DNA extraction and quantification*

DNA was extracted from tissue, scat (both whole scats and swab samples), hair and saliva samples using Qiagen DNeasy kits following the manufacturer’s protocol. For samples collected from the wild, species identity was determined by sequencing a short fragment from the *cyt-b* region (Farrell, Roman, & Sunquist, 2000) and using BLAST to match the sequence. DNA was quantified using a qPCR assay with c-myc primers (Bhavanishankar, Anuradha Reddy, Gour, & Shivaji, 2013) for non-invasive samples, and quantified with qubit for tissue samples. This is the first step presented in Figure 1a, and for the tiger samples resulted in highly varied DNA concentrations (Figure 2a).

*Library Preparation*

Library preparation was essentially a three step process – two PCR reactions followed by pooling. These steps are shown in Figure 1a (multiplex PCR, indexing PCR and pool). The first step was the multiplex PCR, where all target regions were simultaneously amplified for each sample. Template DNA volume was adjusted such that no sample had more than 1 ng per reaction. PCR products from this PCR were diluted 200 times and used as template for the next, indexing PCR where a unique combination of forward and reverse barcodes was assigned to each sample. Equal volumes of all barcoded products were pooled and cleaned in a two-step reaction with Ampure beads. The library was run on a bioanalyzer to test amplification and then sequenced on a Miseq platform (as shown in Figure 1 a, after pool library).

c. Samples chosen for testing

The original primer set targeting 192 SNPs was tested on a panel of captive tiger samples (Table S1). This included samples from captive tiger belonging to three subspecies (Amur, Malayan, Sumatran) and generics (subspecies not known) in the US. This set of samples also included an artificially aged sample from the zoo (to test the effect of sample age) and a serially diluted sample (to test the effect of host DNA concentration). For 5 individuals, both fecal and tissue samples were included in order to compare the fecal genotypes to that obtained from tissues. All samples were genotyped in triplicate to see repeatability (precision) across replicate genotypes.

The 126 primer set was tested primarily on field samples from multiple sources (fecal samples - both swabs and whole feces, hair, saliva swabs and blood) and locations in India, with a wide range of host DNA concentration. The test set samples included the sympatric carnivores leopard and dhole because their feces may be confused with tiger feces. In all cases, samples were typed in triplicate to measure genotyping error across repeats of the same sample (a measure of precision). The sample set included paired tissue and fecal samples from five zoo individuals, of which two individuals shared a parent - offspring relationship and another two were siblings (based on information provided by the zoo). The details of these samples are listed in Table S2.

d. Bioinformatics and analysis

This section refers to the bioinformatics and genotyping steps presented in Figure 1a. Raw reads were first trimmed to remove adapters and bases with quality above 30 using program trimgalore (<https://www.bioinformatics.babraham.ac.uk/projects/trim_galore/>; quality 30, stringency 5, length 5). Trimmed reads were aligned to the reference genome using BWA (Li & Durbin, 2009; bwa mem was used to align)and indexed. SNPs were called from these files using samtools/bcftools (Li et al., 2009; bcftools mpileup was used with the annotate option to allow for subsequent filtering, followed by bcftools call with -c) and the vcf file obtained was filtered to retain only the targeted SNP positions using vcftools(Danecek et al., 2011).

*192 SNP panel*

Amplification success of the 192 primer pairs tested in the first panel was assessed through read counts obtained for each SNP. Genotyping success of each SNP was used to determine whether or not it was included in the second panel. In addition, a trial run was done with all the 192 primers on field samples from India (data not shown), following the same protocol outlined above. Primers that fail to amplify targets nevertheless use up reads. Therefore, primers for SNPs that were successfully typed (genotyped in at least 50% of samples tested) in both runs were chosen for further analysis on field samples, reducing the panel to 126 SNPs.

126 *Panel*
i. SNP filtering

SNP data from the 126 primer set was filtered based on variant quality and missing data. The variant file was filtered to remove genotypes that had read depth or genotype quality lower than 10. Following this, SNPs with over 50% missing genotypes from individuals were filtered out, leaving 114 SNPs (Danecek et al., 2011). From this data, concordance of genotypes for two types of comparisons (within-sample and within individual - between tissue types) and genotyping success for each sample were calculated in R.

ii. Sample filtering

For other analyses, samples with a high proportion of missing data (not typed at 60% or more SNPs) were discarded(Danecek et al., 2011). This data was used to calculate Probability of Identity (P_ID_) (Peakall& Smouse, 2006), to test for recaptures in the wild, assess population structure (Pritchard, Stephens, & Donnelly, 2000).

iii. Calculations/Analyses

Agreement of genotypes across technical replicates of the same sample was calculated for all unique samples. To calculate agreement, the proportion of matching genotypes for each triplicate was recorded and the mean across three comparisons for each sample was calculated in R. Fecal and tissue genotypes were also compared for five individuals, since fecal genotypes are typically considered to be more error prone. Genotyping success was calculated as the proportion of SNPs typed for each sample.

Relatedness was estimated for 77 samples using the PI_HAT estimator (Purcell et al., 2007). Probability of identity (P_ID_) was calculated with only known unique individuals (zoo individuals or wild caught samples). Genotypes were compared across the data to look for recaptures. Genotypes from wild samples (single repeat) were used to assess population structure. The program was run with the admixture and correlated allele frequencies models for 100,000 burn-in steps, followed by 1,000,000 MCMC runs. To select the best model (K), delta K statistic was used, following Evanno’s method(Evanno, Regnaut, & Goudet, 2005).

2. **Tiger genotyping: Supplemental Results**

*2a. The effect of sample DNA concentration on genotype success*

For the 192 SNP data, we plotted the mean number of SNPs successfully genotyped for each sample with the log_10_ transformed tiger DNA concentration for each sample (Fig. S1). Visual inspection of this relationship indicates that there is a positive effect of DNA concentration on genotyping success when DNA concentration is low. However, this effect is absent when DNA concentration is high. To test if this observation was statistically significant we first fitted a generalized linear mixed effects model with genotype success as the response variable and concentration, subspecies, and sample type (tissue, or other) as dependent variables. Concentration was scaled to mean 0 and variation 1. We used a binomial error model. To account for pseudoreplication we included random effects for SNP identity and sample identity. We removed SNPs that had fewer than 10 successful genotypes from the dataset. No dependent variables in this initial model had significant effects on genotyping success.

We then fitted the same model with the addition of a breakpoint variable at concentration <= 0.3 ng/ul. This binary variable indicated that the sample was “low concentration”. We included an interaction between the low concentration variable and actual concentration. A significant interaction would indicate that the effect of concentration changes at or near 0.3 ng/ul. This model found a significant negative effect of low concentration (b=-5.79; p = 3.17e-05; z = -4.161) on genotyping success. Additionally, the interaction between low concentration and actual concentration was significant (b = 452.9481; p = < 2e-16; z = 49.893) indicating that when concentration is less than 0.3 ng/ul there is a positive effect of concentration on genotyping success, but that effect is absent when concentration is higher than 0.3 ng/ul. This model also uncovered some small, but significant, subspecies effects on genotyping success such that Bengal tiger samples had lower genotyping success (see Table S4); and a positive effect on genotyping success when the sample type was tissue. Because the majority of Bengal tiger samples were from the wild we believe that sample quality may be somewhat more degraded than from the other subspecies.

*2b. 192 SNP panel*

After read alignment primer pairs for 42 SNPs failed entirely. Further, the trial run on field samples from India (data not shown) showed amplification for 132 primers. Of these, 6 amplified in less than 50% of samples tested (107 samples in total). 126 SNPs that commonly amplified more than half the tested samples in both the first test and the trial test on field samples were retained for the final analysis.

*2c. 126 SNP panel*

12 SNPs did not amplify for over 50% of samples and were excluded from subsequent analysis. Within population, relatedness estimates were higher than (mean= 0.14,range= 0.0-0.64) between population comparisons (mean=0.009, range=0.0-0.38) as could be expected. Among replicates of a sample, relatedness estimates were nearly 1.0, although for some samples with larger proportion of missing data the estimates were lower (mean=0.97, range=0.67-1.0). No recaptures were found in the data, either based on relatedness estimates or the genotype matches in genalex. The two leopard samples, one from a zoo and one wild, appeared to be recaptures, with nearly identical genotypes (on average, only 3 mismatches in genotypes across all sites compared). The relatedness between the two was also estimated to be very high (average PI_HAT) across all comparisons = 0.96). Genetic differentiation was detected among the wild samples. The Delta K value was lowest for K=2 (137.7), followed very closely by K=3 (113.8), hence results for both have been plotted (Fig. S2).

3. **Conch Genotyping: Materials and Methods**

3a. Conch transcriptome:

*Sample Collection*

Queen conch were sampled from six sites throughout their native range: Aruba in the Caribbean Netherlands; Exuma Cays, Bahamas; Carrie Bow Cay, Belize; Florida Keys, USA; Pedro Bank, Jamaica; and St. Eustatius, Caribbean Netherlands (Table S3). A total of 96 adult queen conch were collected by hand (16 from each location), and a 5 mm piece of mantle tissue was collected from each conch using sterilized biopsy forceps. Mantle tissue was placed into a 2 mL cryotube containing an RNA stabilizing reagent and stored at 4^o^ C for 1 day to allow the reagent to penetrate the mantle tissue, then moved to -20^o^ C for 2-4 weeks, and then archived at -80^o^ C.

*Extraction and Sequencing*

We extracted RNA from 96 *L. gigas*individuals using the Direct-zol RNA MiniPrep Kit (Zymo Research) following the manufacturer’s protocol. For each extraction approximately 2 mm^2^ of mantle tissue was sliced away from the larger piece of archived tissue using a sterile razor blade. The mantle tissue was placed in 2 mL Eppendorf tube with an o-ring cap containing 2.3 mm diameter zirconia-silicon beads then homogenized in TRI Reagent for 1minute using Qiagen Tissue lyser. Total RNA was quantified using a Qubit 2.0 florometer (Invitrogen). Extracted RNA was stored at -80 ^o^C for 1-7 days prior to cDNA library preparation.

The TruSeq Stranded mRNA Library Prep Kit (Illumina) was used to create an individually barcoded transcriptome library for each individual. A total of 2 ug of RNA was used to generate each cDNA library using the TruSeq Stranded mRNA sample kit from Illumina (San Diego, CA, USA). We followed the Illumina protocol and one of 16 unique Illumina adaptors were used to demultiplex each sample. The quality of each cDNA library was validated with a Bioanalyzer (Agilent, Santa Clara, CA, USA) and samples were pooled into 6 lanes (16 libraries per lane) of sequencing on an Illumina HiSeq 2000 sequencer. Each lane was sequenced with 125 cycle paired-end reads at the Huntsman Cancer Institute, University of Utah.

*Sequence quality control and Transcriptome assembly*

Processing of raw reads was performed following an updated version of Simple Fools Guide to RNA-Seq (SFG) pipeline (De Wit *et al.* 2012) available at (<https://github.com/bethsheets/Population-Genomics-via-RNAseq>). FastQC v0.11.2 (<http://www.bioinformatics.babraham.ac.uk/projects/fastqc/>) was used to generate reports containing summary statistics and quality assessments of the raw reads. Reads were processed by Trimmomatic v.0.33 (Bolger, Lohse, & Usadel, 2014) to remove left over adaptors from the sequencing process and remove reads with a Phred quality score of below 30 (i.e. >99.9% base call accuracy). After filtering, reads from four queen conch individuals (one individual each from Aruba, Belize, Florida, and St. Eustatius) were imported into TRINITY v2.2.0 (Grabherr et al., 2013) to assemble a *de novo* transcriptome. The parameters followed the default settings recommended by the authors.

The BLASTX algorithm using NCBI’s default parameters was used to identify contigs from the *de novo* transcriptome assembly that matched gene models derived from the genome of *Lottiagigantea*, a marine gastropod mollusc. Matches were considered significant at e- values of 10^-4^. Due to the lack of genomic resources for *Lobatusgigas*, contigs that did not significantly match *Lottiagigantea* gene models were not retained in the final transcriptome assembly. This methodology was used to prevent contaminant sequences from bacteria or unicellular eukaryotes potentially residing within of the mantle tissue from being erroneously included in the *de novo* transcriptome assembly.

*Mapping and SNP calling*

Detection of single nucleotide polymorphisms (SNPs) from functional genomic regions were identified using updated scripts of the SFG pipeline (<https://github.com/bethsheets/Population-Genomics-via-RNAseq>). The paired-end reads from each individual were mapped to the *de novo* transcriptome assembly with BOWTIE 2 (Langmead & Salzberg, 2012) using the very strict default mapping parameters modified to allow only one mismatch. The Bayesian genetic variant detector FREEBAYES v0.9.10 (https://github.com/ekg/freebayes) was then used to detect SNPs from the mapped reads following the default SNP calling parameters.The program package VCFtools(Danecek *et al.* 2011) was used to filter the SNPs identified by FREEBAYES. First, all SNPs with > 2 alleles, and with a minor allele frequency < 0.5% were removed. The SNPs passing the first filter that were present in all individuals and had quality scores (Phred) > 30 and had high coverage > 10X, were retained for population genomic, local adaptation, and assignment testing.

3b. SNP Primer design and testing

We used our G4C (Genotyping for Conservation) script library to design 192 primer pairs with the same traits as for tigers above. We used these primers to genotype 279 conch from 14 populations including 12 samples from conch fritters purchased at restaurants in Miami.93 conch primers produced no data, and another 36 produced genotypes for less than 2/3 of the samples. This low success was most likely because the primers were based on transcriptome data that cannot discern intron-exon boundaries (Fig. S3).

Each row represents genotype data from one sample. Each column represents one of 62 SNP loci. Red colors indicate missing data, for fresh conch samples (first set, shown in order of decreasing success rate) and samples of fried conch.

4. **Conch Genotyping: Supplemental Results**

***Transcriptome SNPs and genetic structure***

Transcriptome sequencing resulted in an average of 21.7 million (m) paired-end reads per individual for a total of 2.08 billion reads. The number of reads per individual after quality trimming adaptors and removing reads with a quality score below 30 ranged from 20.5 m – 5.2 m reads per individual. The initial *de novo* transcriptome assembly contained 563,514 contigs. After quality control and retaining only contigs that aligned to gene models from the *Lottia gigantean* genome, the final filtered *de novo* transcriptomeassembly consisted of 55,712 contigs with a mean length of 1,261 base-pairs (median 696 base-pairs). The percentage of contigs < 300 base pairs was 19.6%. A total of 42,780 (76.8%) contigs were annotated in the final assembly.

The preliminary SNP detection using Freebayes identified a total of 480,962 SNPs with a Phred quality score > 30. These SNPs were filtered to retain biallelic SNPs with a minor allele frequency > 0.5%, with high coverage > 10X, Phred quality scores > 30, and present in all individuals with no missing data. The resulting panel of SNPs contained total of 14,841 SNPs. This filtered panel of SNPs was retained for population genomic and traceability analyses in a total of 83 individuals.

**Population Structure**

The panel of 14,297 SNP sidentified significant pairwise levels of population differentiation, *F*_ST_ (Table S5). A total of 6 out of 15 pairwise combinations of *F*_ST_ were significant (False Discovery Rate (FDR) adjusted *p* < 0.008) across the six sampling locations that represent a selection of *L*. *gigas*fisheries regulated by CITES. Levels of differentiation were the strongest between Aruba & Bahamas (pairwise *F*_ST_ = 0.012). Pairwise levels of population differentiation were significant for 6 out of 15 pairwise combinations (False Discovery Rate adjusted *p* < 0.0041). The global *F*_ST_ across the six sampling locations was 0.0046.

**GTSeq comparisons**

*Genetic identity and close kin:* Euclidean genetic distance among conch samples (average 7.76, Fig. S4, calculated using scripts from https://popgen.nescent.org/2015-05-18-Dist-SNP.html) shows only a single peak with no secondary peaks where close kin such as parent-offspring pairs or full or half sibs should occur (Fig. S5). Two pairs of individuals showed genetic identity: One of these was a duplicated control sample from St. Eustatius (Fig. S5). A second control duplicate from Sand Bores cay showed a genetic distance of 1.6, corresponding to a discrepancy at a single locus. All five conch samples with genetic distances between 0-1.6 derive from control duplicates (n=3) or conch sampled from the same fishing boat (n=2).

Supplementary Figures:

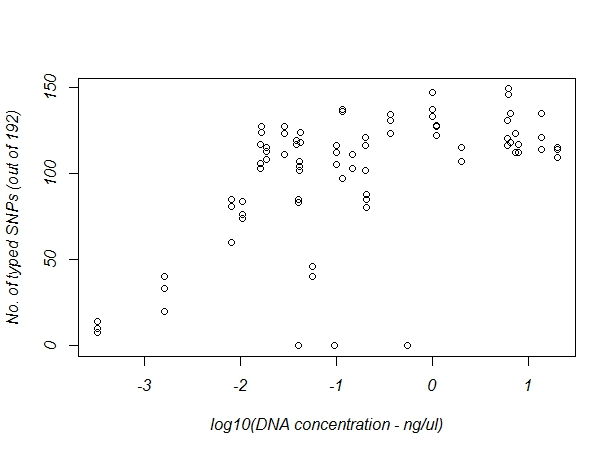

Figure S1: Number of SNPs typed for varying DNA concentration from the 192 SNP panel data.

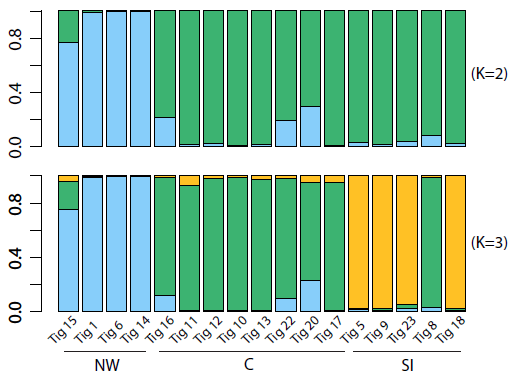

Figure S2: Structure plot depicting genetic clusters identified in the wild tiger samples. The highest support based on deltaK was for two genetic clusters in the data (upper panel), followed by three clusters (lower panel). The map on the right depicts the sampling locations. The location names indicate geographically clustered regions (NW = North-West, N = North, C = Central India, SI = South India).

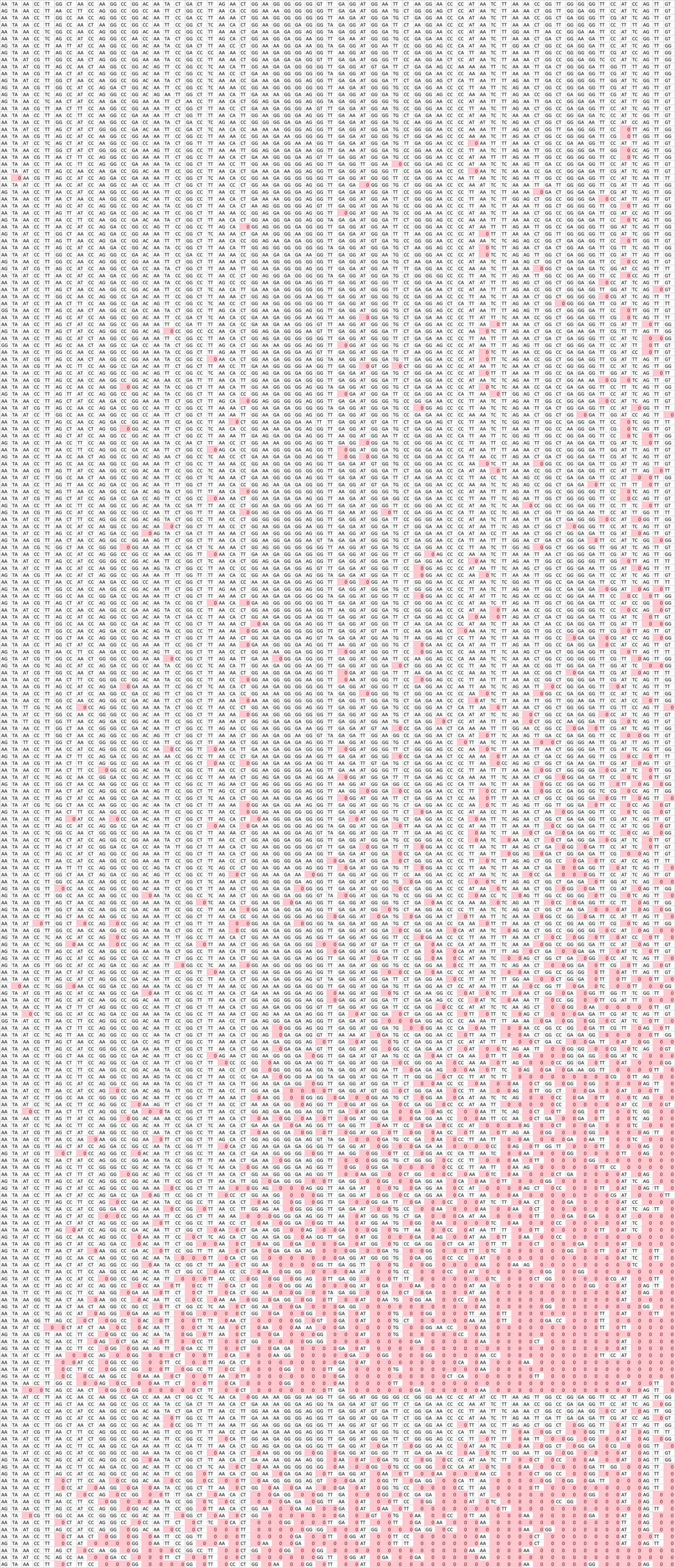

Figure S3: Genotype success for 199 conch samples and 25 samples isolated from fried conch. Individuals are in rows. SNPs are arrayed in columns. Red squares denote lack of genotype data.

Figure S4: Full pairwise Euclidean distance matrix for 185 conch samples including 12 from fried fritters. Cells are colored proportional to genetic similarity (red = 1; green = 0). The diagonal compares each individual to itself.

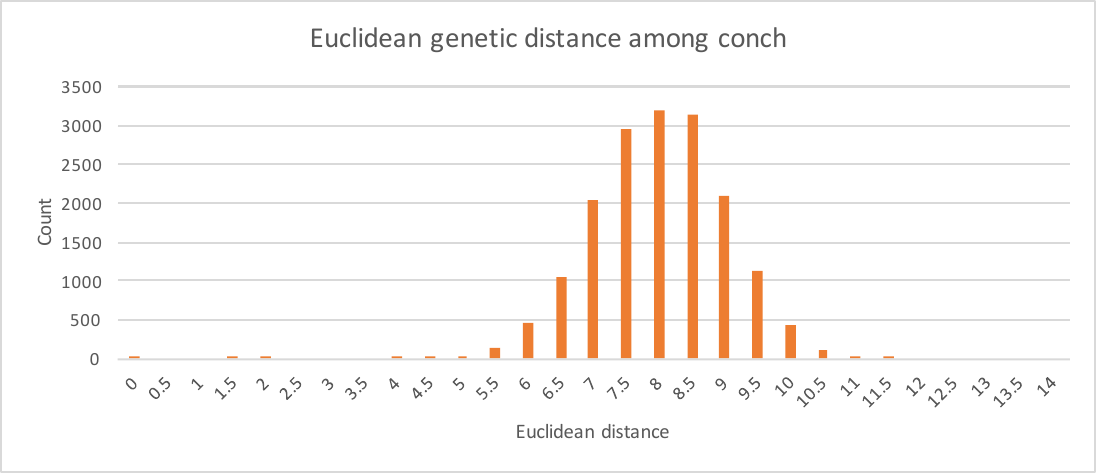

Figure S5: Distribution of Euclidean distances among conch all samples.

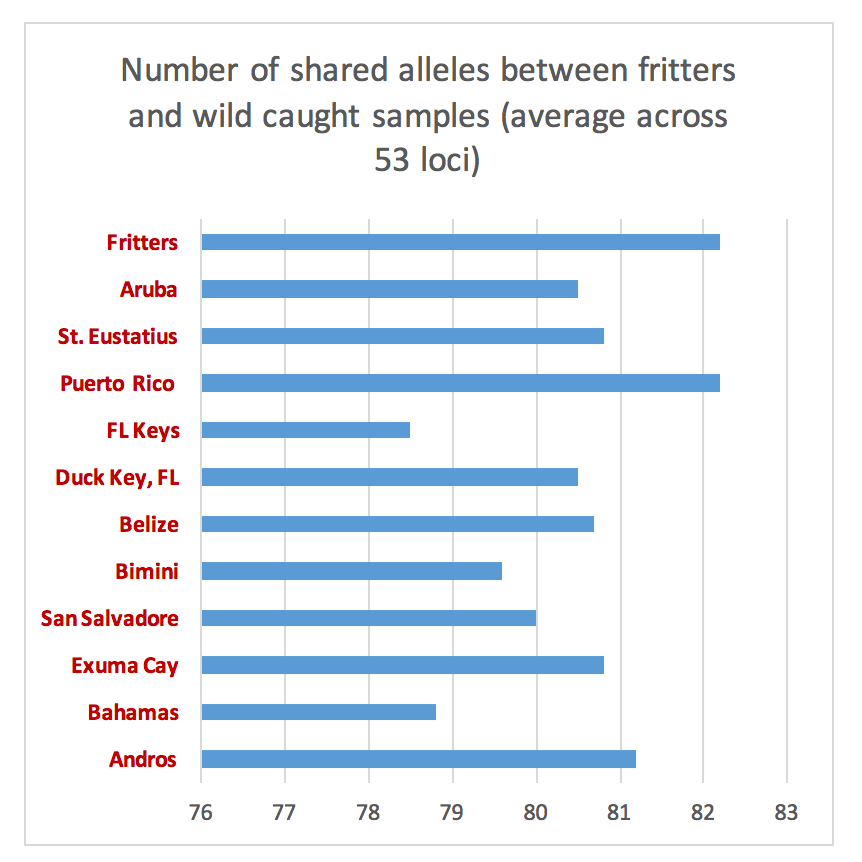

Figure S6: Average genetic similarity among conch sampled from commercial fritters compared to populations throughout the Caribbean. Average relative genetic identity between fritter samples and conch from the Florida Keys is significantly lower (p<0.001, binomial test) than compared to other populations (Fig. S6). The highest similarities between fritters and sampled populations are for Puerto Rico, St. Eustatius and Andros Island. The lowest are for the Florida Keys and Nassau.

Supplementary Tables:

| Sample ID | Sample type | Concentration(ng/ul) | Subspecies |
| --- | --- | --- | --- |
| Amur 1 | Fecal | 0.0285 | Amur |
| Amur 1 | Tissue | 7.4300 | Amur |
| Amur 2 | Fecal | 0.0164 | Amur |
| Amur 2 | Fecal | 0.0945 | Amur |
| Amur 3 | Tissue | 2.0200 | Amur |
| Amur 4^#^ | Tissue | 1.0000 | Amur |
| Amur 4^#^ | Tissue | 0.2000 | Amur |
| Amur 4^#^ | Tissue | 0.0400 | Amur |
| Amur 4^#^ | Tissue | 0.0080 | Amur |
| Amur 4^#^ | Tissue | 0.0016 | Amur |
| Amur 4^#^ | Tissue | 0.0003 | Amur |
| Blank | Blank | 0.0000 | Blank |
| Generic 1 | Tissue | 6.1400 | Generic |
| Generic 1^$2^ | Fecal | 6.1530 | Generic |
| Generic 1^$2^ | Fecal | 0.3645 | Generic |
| Generic 1^$2^ | Fecal | 0.1475 | Generic |
| Generic 1^$1^ | Fecal | 1.1115 | Generic |
| Generic 1^$3^ | Fecal | 0.0163 | Generic |
| Generic 1^$3^ | Fecal | 0.0184 | Generic |
| Generic 2 | Fecal | 0.5539 | Generic |
| Generic 2 | Tissue | 13.6000 | Generic |
| Generic 3 | Fecal | 0.0560 | Generic |
| Generic 3 | Tissue | 20.0400 | Generic |
| Generic 4 | Fecal | 0.2065 | Generic |
| Generic 5 | Fecal | 0.1145 | Generic |
| Generic 5* | Fecal | 0.1145 | Generic |
| Malayan 1 | Fecal | 0.0104 | Malayan |
| Malayan 1 | Fecal | 0.1002 | Malayan |
| Malayan 2 | Fecal | 0.0379 | Malayan |
| Malayan 2 | Fecal | 0.0419 | Malayan |
| Sumatran 1 | Tissue | 6.5200 | Sumatran |
| Sumatran 2 | Tissue | 7.8500 | Sumatran |
| Sumatran 2 | Fecal | 0.0413 | Sumatran |

Table S1: List of samples chosen for the first test with 192 primer pairs. This includes a serially diluted sample (#), an artificially aged sample ($1 – collected within 2-3 hours, $2 – within 11 hours, $3 – within 48 hours). * indicates higher input volume of sample (2 ul).

| Sample | DNA (ng/ul) | Type | Location | Region | Species |
| --- | --- | --- | --- | --- | --- |
| Blank | 0.000 | Water |  |  | blank |
| Dhole.1 | 1.920 | Tissue | Zoo | ZOO | Dhole |
| Leopard.1 | 10.700 | Tissue | Zoo | ZOO | Leopard |
| Leopard.2 | 0.179 | feces (swab) | Chandrapur | CI | Leopard |
| Tiger.1 | 22.500 | Tissue | Ranthambore | NW | Tiger |
| Tiger.10 | 3.286 | feces (swab) | Zoo | ZOO | Tiger |
| Tiger.11 | 1.288 | feces (swab) | Zoo | ZOO | Tiger |
| Tiger.12 | 0.832 | Tissue | Periyar | SI | Tiger |
| Tiger.13 | 0.405 | feces (swab) | Nagarahole | SI | Tiger |
| Tiger.14 | 0.276 | Saliva | Kanha | CI | Tiger |
| Tiger.15 | 0.159 | Shed Hair | Kanha | CI | Tiger |
| Tiger.16 | 0.139 | Saliva | Kanha | CI | Tiger |
| Tiger.17 | 0.072 | Shed Hair | Kanha | CI | Tiger |
| Tiger.18 | 0.064 | feces (swab) | Ranthambore | NW | Tiger |
| Tiger.19 | 0.063 | feces (swab) | Ranthambore | NW | Tiger |
| Tiger.2 | 21.400 | Tissue | Zoo | ZOO | Tiger |
| Tiger.20 | 0.055 | feces (swab) | Zoo | ZOO | Tiger |
| Tiger.21 | 0.050 | feces (whole) | Corbett | N | Tiger |
| Tiger.22 | 0.016 | feces (whole) | Kanha-Pench | CI | Tiger |
| Tiger.23 | 0.015 | feces (swab) | Wayanad | SI | Tiger |
| Tiger.24 | 0.007 | feces (swab) | Pench | CI | Tiger |
| Tiger.25 | 0.005 | feces (swab) | Brahmapuri | CI | Tiger |
| Tiger.26 | 0.005 | feces (swab) | Ranthambore | NW | Tiger |
| Tiger.27 | 0.002 | feces (whole) | Bandhavgarh | CI | Tiger |
| Tiger.28 | 0.002 | feces (swab) | Nagarahole | SI | Tiger |
| Tiger.3 | 20.600 | Tissue | Zoo | ZOO | Tiger |
| Tiger.4 | 13.100 | Tissue | Zoo | ZOO | Tiger |
| Tiger.5 | 8.865 | feces (swab) | Zoo | SI | Tiger |
| Tiger.6 | 8.692 | feces (swab) | Zoo | ZOO | Tiger |
| Tiger.7 | 7.833 | feces (swab) | Ranthambore | NW | Tiger |
| Tiger.8 | 6.100 | Tissue | Zoo | ZOO | Tiger |
| Tiger.9 | 5.800 | Tissue | Zoo | SI | Tiger |

Table S2: List of samples chosen for the final test with 126 primer pairs. The locations NW, N, CI, SI refer to Northwest, North, Central and Southern India.

| Site | Country | *N* | Lat | Long | Date |
| --- | --- | --- | --- | --- | --- |
| Aruba | Caribbean Netherlands | 16 | 12.417 | -69.889 | Feb-16 |
| Carrie Bow Caye | Belize | 16 | 16.803 | -88.082 | Aug-15 |
| Exuma Cayes | Bahamas | 16 | 24.337 | -76.597 | Jun-15 |
| Florida Keys | USA | 16 | 24.743 | -80.824 | Jun-15 |
| Pedro Bank | Jamaica | 16 | 16.945 | -78.720 | Jul-15 |
| Saint Eustatius | Caribbean Netherlands | 16 | 17.480 | -62.989 | Sep-15 |

**Table S3.** Sample collection information for Queen conch transcriptome analysis.

|  | Estimate | Std. Error | z value | Pr(>\|z\|) |
| --- | --- | --- | --- | --- |
| (Intercept) | -1.6807 | 1.6464 | -1.021 | 0.3073 |
| concentration | -0.0124 | 0.1729 | -0.072 | 0.9428 |
| low_concentrationTRUE | -5.7901 | 1.3915 | -4.161 | 3.17e-05 *** |
| subspecies_generic | 2.9531 | 1.7432 | 1.694 | 0.0903 . |
| subspecies_malayan | 4.6397 | 2.1302 | 2.178 | 0.0294 * |
| subspecies_sumatran | 2.6523 | 2.0998 | 1.263 | 0.2065 |
| type_tissue | 3.6985 | 1.7668 | 2.093 | 0.0363 * |
| concentration:low_concentrationTRUE | 452.9481 | 9.0785 | 49.893 | < 2e-16 *** |
| Random effects:  Groups Name Variance Std.Dev.  snp (Intercept) 8.361 2.892  sample (Intercept) 9.928 3.151  Number of obs: 12320, groups: snp, 138; sample, 32 | | | | |

Table S4: Estimates from mixed effects model on the 192 SNP panel.

| Transcriptome SNPs | Aruba | Bahamas | Belize | Florida | Jamaica | St. Eustatius |
| --- | --- | --- | --- | --- | --- | --- |
| Aruba | __ | **0.0124** | **0.0065** | **0.0033** | **0.0054** | 0.0019 |
| Bahamas | **0.0124** | __ | 0.0074 | 0.0048 | 0.0059 | **0.0088** |
| Belize | **0.0065** | 0.0074 | __ | 0.0036 | 0.0053 | 0.0040 |
| Florida | **0.0033** | 0.0048 | 0.0036 | __ | 0.0010 | **0.0023** |
| Jamaica | **0.0054** | 0.0059 | 0.0053 | 0.0010 | __ | 0.0008 |
| St. Eustatius | 0.0019 | **0.0088** | 0.0040 | **0.0023** | 0.0008 | __ |

**Table S5.** Pairwise *F*_ST_ for a panel of 14,297 putatively neutral transcriptome derived single nucleotide polymorphisms from Queen conch.

References

Bhavanishankar, M., Anuradha Reddy, P., Gour, D. S., & Shivaji, S. (2013). Validation of non-invasive genetic identification of two elusive, sympatric, sister-species- tiger (panthera tigris) and leopard (panthera pardus). *Current Science*, *104*(8), 1063–1067.

Bolger, A. M., Lohse, M., & Usadel, B. (2014). Trimmomatic: A flexible trimmer for Illumina sequence data. *Bioinformatics*, *30*(15), 2114–2120. doi:10.1093/bioinformatics/btu170

Campbell, N. R., Harmon, S., & Narum, S. R. (2014). Genotyping-in-Thousands by sequencing (GT-seq): A cost effective SNP genotyping method based on custom amplicon sequencing. *Molecular Ecology Resources*, n/a-n/a. doi:10.1111/1755-0998.12357

Danecek, P., Auton, A., Abecasis, G., Albers, C. a., Banks, E., DePristo, M. a., … Durbin, R. (2011). The variant call format and VCFtools. *Bioinformatics*, *27*(15), 2156–2158. doi:10.1093/bioinformatics/btr330

Evanno, G., Regnaut, S., & Goudet, J. (2005). Detecting the number of clusters of individuals using the software STRUCTURE: a simulation study. *Molecular Ecology*, *14*(8), 2611–20. doi:10.1111/j.1365-294X.2005.02553.x

Farrell, L. E., Roman, J., & Sunquist, M. E. (2000). Dietary separation of sympatric carnivores identified by molecular analysis of scats. *Molecular Ecology*, *9*(10), 1583–90. Retrieved from http://www.ncbi.nlm.nih.gov/pubmed/11050553

Garrison, E., & Marth, G. (2012). Haplotype-based variant detection from short-read sequencing. *ArXiv Preprint ArXiv:1207.3907 [q-Bio.GN]*, 9. doi:arXiv:1207.3907 [q-bio.GN]

Grabherr, M. G., Haas, B. J., Yassour, M., Levin, J. Z., Thompson, D. A., Amit, I., … W., B. (2013). Trinity: reconstructing a full-length transcriptome without a genome from RNA-Seq data. *Nature Biotechnology*, *29*(7), 644–652. doi:10.1038/nbt.1883.Trinity

Langmead, B., & Salzberg, S. L. (2012). Fast gapped-read alignment with Bowtie 2. *Nature Methods*, *9*(4), 357–359. doi:10.1038/nmeth.1923

Li, H., & Durbin, R. (2009). Fast and accurate short read alignment with Burrows–Wheeler transform. *Bioinformatics*, *25*(14), 1754–1760. Retrieved from http://dx.doi.org/10.1093/bioinformatics/btp324

Li, H., Handsaker, B., Wysoker, A., Fennell, T., Ruan, J., Homer, N., … Durbin, R. (2009). The Sequence Alignment/Map format and SAMtools. *Bioinformatics*, *25*(16), 2078–2079. doi:10.1093/bioinformatics/btp352

Peakall, R., & Smouse, P. E. (2006). GENALEX 6: genetic analysis in Excel. Population genetic software for teaching and research. *Molecular Ecology Notes*, *6*, 288–295.

Pritchard, J. K., Stephens, M., & Donnelly, P. (2000). Inference of population structure using multilocus genotype data. *Genetics*, *155*(2), 945–959.

Purcell, S., Neale, B., Todd-Brown, K., Thomas, L., Ferreira, M. a R., Bender, D., … Sham, P. C. (2007). PLINK: a tool set for whole-genome association and population-based linkage analyses. *American Journal of Human Genetics*, *81*(3), 559–75. doi:10.1086/519795
